# Comparative Study of High-resolution LysB29(N_ε_-myristoyl) des(B30) Insulin Structures Display Novel Dynamic Causal Interrelations in Monomeric-Dimeric Motions

**DOI:** 10.1101/2022.11.19.517203

**Authors:** Esra Ayan, Ebru Destan, Abdullah Kepceoğlu, Halilibrahim Ciftci, Ahmet Katı, Hasan DeMirci

## Abstract

The treatment of insulin-dependent diabetes mellitus is characterized by artificial supplementation of pancreatic β-cell ability to regulate sugar levels in the blood. Even though various insulin analogs are crucial for reasonable glycemic control, understanding the dynamic mechanism of the insulin analogs may help to improve the best-protracted insulin analog to assist people with Type 1 Diabetes (T1D) to live comfortably while maintaining tight glycemic control. Here we present the high-resolution crystal structure of NN304, known as insulin detemir, to 1.7 Å resolution at cryogenic temperature. We computationally further investigated our crystal structure’s monomeric-dimeric conformation and dynamic profile by comparing it with a previously available detemir structure (PDB ID: 1XDA). Our structure (PDB ID: 8HGZ) obtained at elevated pH provides electrostatically triggered minor movements in the equilibrium between alternate conformational substates compared to the previous structure, suggesting it might induce an intermediate state in the dissociation pathway of the insulin detemir’s hexamer:dihexamer equilibrium. Supplemented with orientational cross-correlation analysis by Gaussian Network Model (GNM), this alternate oligomeric conformation offers the distinct cooperative motions originated by loose coupling of distant conformational substates of a protracted insulin analog that has not been previously observed.

## Introduction

In recent years the number of individuals with diabetes has increased at a higher rate in low and middle-income countries than in high-income countries (WHO). Additionally, in 2019, diabetes was recorded as the ninth leading cause of death, considering a predicted 1.5 million deaths directly caused by diabetes (WHO). Pancreatic insulin delivery can be challenging to reproduce by subcutaneous injection [1]. Among individuals with both Type 1 Diabetes (T1D) and Type 2 Diabetes (T2D), the majority of the patients fail to manage glycemic targets resulting in an increased risk of various complications [2]. The primary purpose of insulin therapy relies on the formulation of both long-acting and rapid-acting insulin analogs. Insulin therapy mimics the function of physiological insulin secretion levels to control endogenous hepatic glucose levels in the blood [3]. Recombinant insulin is a critical peptide hormone for treating insulin-dependent diabetes mellitus (DM) [3,4]. During the last decade, insulin preparations have been significantly modified to reduce the risk of health complications due to breakthroughs in the understanding of insulin structure and its complex aggregation properties [5]. The efficiency, efficacy and simplicity of insulin analogs can facilitate the patient’s transition to insulin therapy [6,7] (Figure S1). Various insulin analogs provide advantages consisting of greater convenience (Figure S1), improved physiologic profile and a reduced risk of hypoglycemia [8]. It is of considerable importance that insulin analogs are long-acting, predictable, and have a flat pharmacodynamic profile to limit the risk of hypoglycemia in diabetic patients [8].

The active form of insulin circulates in the blood as a monomer protein consisting of two polypeptide chains: an A chain of 21 amino acids and a B chain of 30 amino acids. Insulin is produced by pancreatic β-cells as a monomer and stored in a hexameric form. The hexameric form is generated by three insulin dimers aggregating around two central zinc cations [7]. Various insulin analogs exist in the market which have been genetically engineered and formulations characterized by residual substitutions over the recombinant DNA technology [9]. Over the years, the commonly known strategy to produce a long-acting insulin variant is to formulate amorphous suspensions or crystalline insulin that provide a slowly dissolvable depot by subcutaneous injection [10].

The LysB29-tetradecanoyl, des-(B30) human insulin (NN304, insulin detemir) was introduced to the market with the brand name Levemir® in the mid-late 90s [11]. Detemir has been created as a neutral, soluble insulin preparation by covalently linking a hydrophobic myristate fatty acid chain to Lys 29 in chain B which is not strictly essential for the activity of human insulin [11]. The unique mechanism of this covalently modified insulin analog relies on its reversible binding to human albumin through the fatty acid extension [6,11]. This albumin bound form of insulin serves as a reservoir that continually and slowly releases detemir insulin into the bloodstream and delays the hormone’s action [6,11,12] (Figure S2). Detemir has demonstrated greater efficacy and reproducible absorption rate compared to neutral protamine hagedorn (NPH) insulin and insulin glargine [13,14,15]. This is why detemir has had a favorable profile in clinical usage for over a decade.

Collection of studies have been conducted on detemir to understand its structure, function, and its interaction with serum albumin [12,16]. In this work, we present the high resolution detemir crystal structure determined to 1.7 Å resolution at the cryogenic temperature. Diffraction data collected at the Turkish Light Source (AKA Turkish DeLight) [17]. Additionally, we employed computational Gaussian Network Model (GNM) analysis that we aimed at investigating the differences of monomer:dimer equilibrium dynamics between our high-resolution cryogenic structure with previously published detemir structure (PDB ID: 1XDA). The crystallographic data presented here provide a distinct novel unit cell origin in asymmetric unit compared to the previously published detemir unit cell orientation [11]. GNM analysis performed on the high-resolution structure also displays a previously unobserved pattern of monomeric-dimeric correlated motion of this long-acting insulin analog. Understanding monomer-dimer correlated motion of the detemir structure still remains to have immense importance in revealing the structural dynamics of the detemir and pharmacokinetics of this important hormone. This will pave the way for improving, modulating, and optimizing the protracted dissociation rate of the hexameric form of detemir into its active monomeric form.

## Results

Throughout the analysis of the manuscript, the renumbered version of 8HGZ and 1XDA structures were used and uploaded as a supplementary file (File S1) under the name Renumbered_8HGZ.pdb, and Renumbered_1XDA.pdb, respectively.

### Crystal Structure of Novel Detemir Crystal Form

Insulin has two peptide chains: the A chain and the B chain. Two disulfide bonds link A and B chains, known as one “monomer.” If two monomers bind each other noncovalently, it’s known as a dimer. The inactive and extracellular released form of insulin is always one hexamer. After the dissociation of the insulin to its monomer form, the hormone is active and can bind to its receptor properly. Detemir insulin is a different analog in terms of its structural hierarchy than its counterparts’ analogs. It includes two hexamers in the unit cell due to the presence of the myristic acids [7]. Therefore, the detemir’s asymmetric unit is also different from other analogs due to the dimer of dimer conformation. Accordingly, the monomer form of the detemir’s dimer of dimer form refers to chains A and B together (or C and D, or E and F, or G and H), and each dimer refers to the pair AB-CD, and EF-GH respectively.

In this study, the cryogenic detemir structure determined to 1.7□Å resolution contains four nearly identical monomer molecules in the asymmetric unit cell, as well as four myristic acid side chains, four phenol molecules, four zinc ions, four chloride ions, and 188 water molecules (Figure 1; Figure S3; FigureS 4; Table 1).

**Figure 1.**
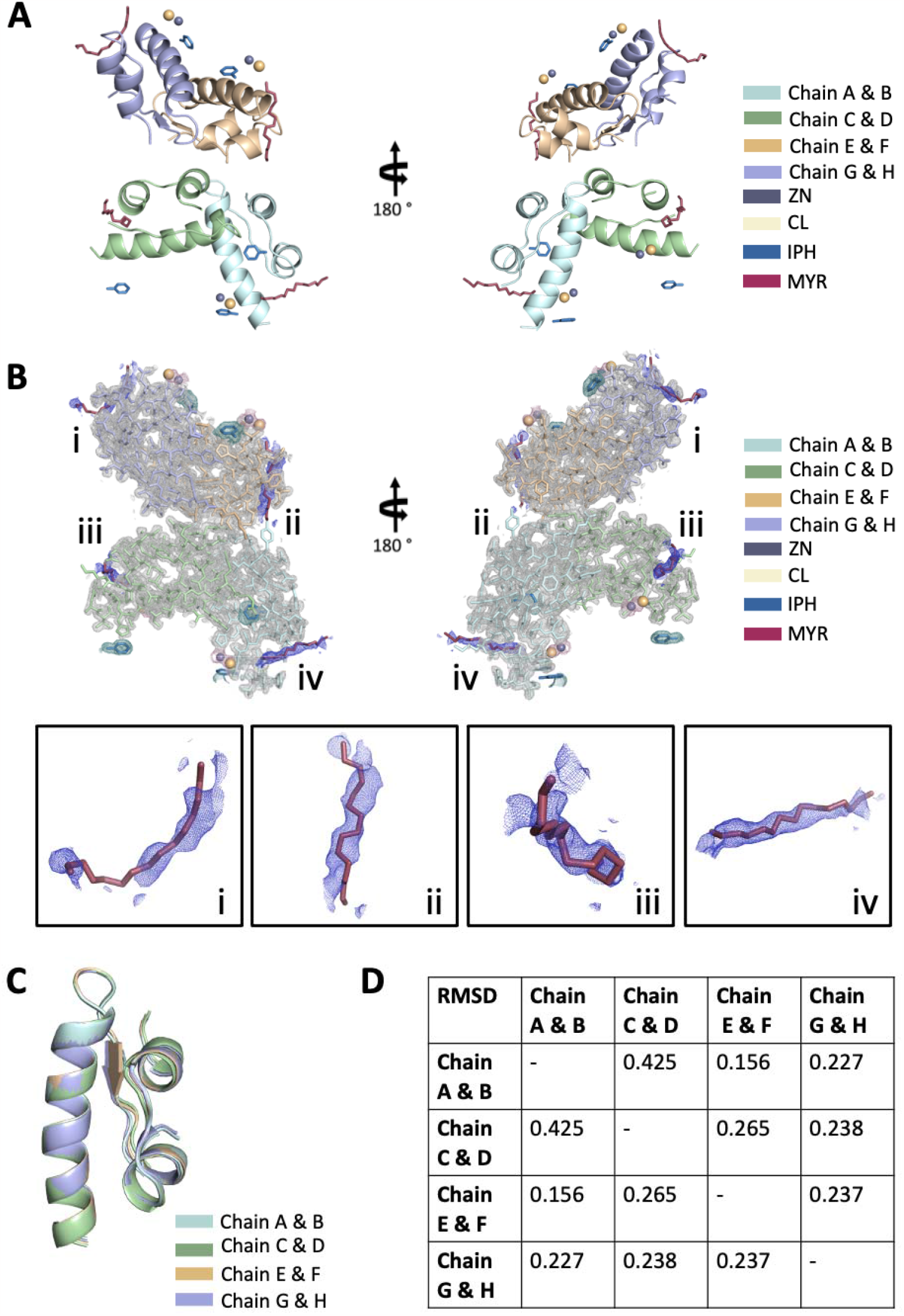
The 8HGZ structure at cryogenic temperature. (A) Detemir structure is represented in cartoon view. Myristic acid (MYR), zinc (ZN), chloride (CL), and phenol (IPH) are shown in stick representation. (B) The refined 2Fo-Fc electron density map covering detemir is shown in gray, electron density for MYR is colored in blue, ZN and CL are colored in dark salmon, and IPH is colored in smudge. (C) Each monomer of detemir is superposed in PyMOL, and (D) RMSD values are indicated in the table.

**Table 1.**
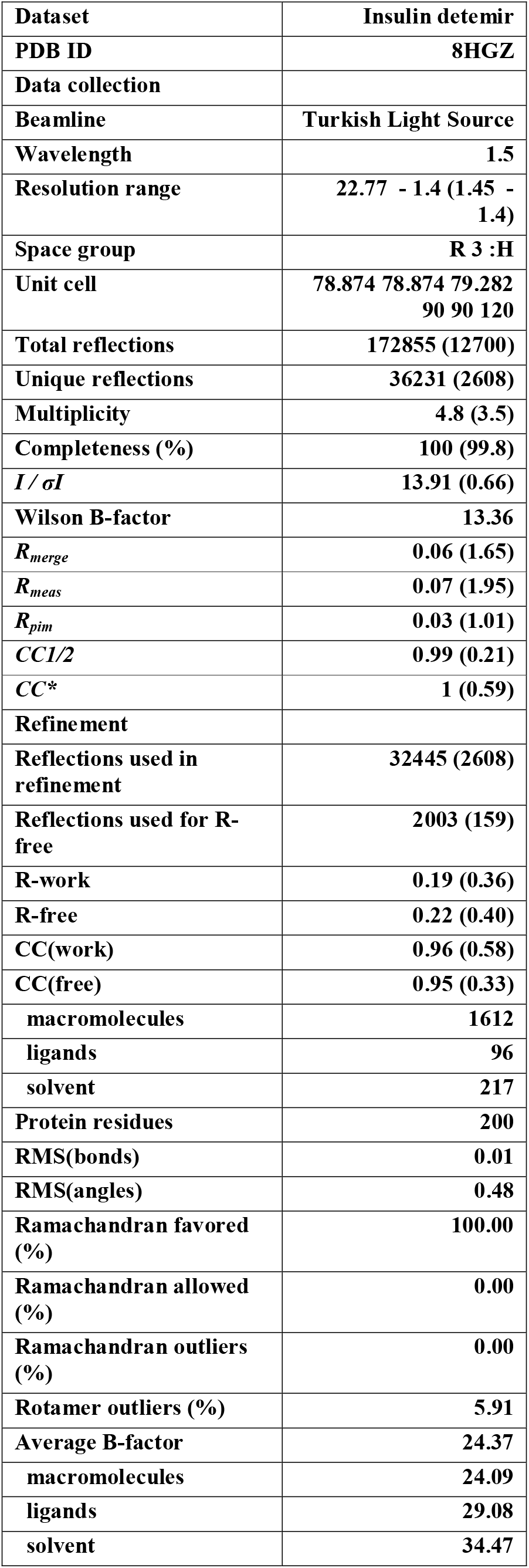
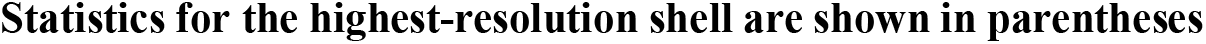
Data collection and refinement statistics.

|  |  |
| --- | --- |
| **Dataset** | **Insulin detemir** |
| **PDB ID** | **8HGZ** |
| **Data collection** |  |
| **Beamline** | **Turkish Light Source** |
| **Wavelength** | **1.5** |
| **Resolution range** | **22.77 - 1.4 (1.45 - 1.4)** |
| **Space group** | **R 3 :H** |
| **Unit cell** | **78.874 78.874 79.282**<br>**90 90 120** |
| **Total reflections** | **172855 (12700)** |
| **Unique reflections** | **36231 (2608)** |
| **Multiplicity** | **4.8 (3.5)** |
| **Completeness (%)** | **100 (99.8)** |
| **$I / \sigma I$** | **13.91 (0.66)** |
| **Wilson B-factor** | **13.36** |
| **$R_{merge}$** | **0.06 (1.65)** |
| **$R_{meas}$** | **0.07 (1.95)** |
| **$R_{pim}$** | **0.03 (1.01)** |
| **$CC1/2$** | **0.99 (0.21)** |
| **$CC^*$** | **1 (0.59)** |
| **Refinement** |  |
| **Reflections used in refinement** | **32445 (2608)** |
| **Reflections used for R-free** | **2003 (159)** |
| **R-work** | **0.19 (0.36)** |
| **R-free** | **0.22 (0.40)** |
| **CC(work)** | **0.96 (0.58)** |
| **CC(free)** | **0.95 (0.33)** |
| **macromolecules** | **1612** |
| **ligands** | **96** |
| **solvent** | **217** |
| **Protein residues** | **200** |
| **RMS(bonds)** | **0.01** |
| **RMS(angles)** | **0.48** |
| **Ramachandran favored (%)** | **100.00** |
| **Ramachandran allowed (%)** | **0.00** |
| **Ramachandran outliers (%)** | **0.00** |
| **Rotamer outliers (%)** | **5.91** |
| **Average B-factor** | **24.37** |
| **macromolecules** | **24.09** |
| **ligands** | **29.08** |
| **solvent** | **34.47** |

Statistics for the highest-resolution shell are shown in parentheses

Our dimer of dimer crystal structures at cryogenic temperature could not be superimposed with the dimer of dimer 1XDA structure due to the distinct crystal packing of the detemir in the asymmetric unit cells (Figure 2; Figure S5; Figure S6). The two dimers of the 1XDA structure form introverted monomers packed side by side, while one of the 8HGZ structure’s dimers displays extroverted monomers in the asymmetric unit cell. Each disulfide bonded monomer chains are labeled as AB, CD, EF, and GH (Figure 1A, B; Figure S3A). Each extroverted monomer in dimer displays more plasticity/flexibility compared to the body/center of the four-monomer structure in the ellipsoid representation which is derived from TLS analysis on PyMOL (Figure S3B). The 8HGZ structure also displays more negative charge distribution in the structure’s central region compared to its periphery (Figure S3C). Each monomer includes a 14-carbon fatty acid chain covalently conjugated to the B29 Lys residue. The myristic acids engage in the crystal contacts among the neighboring dimers, radiating out the aliphatic side chains from the crystallographic axis toward the periphery (Figure S4). The “dimer of dimer” asymmetric form of 8HGZ shows minor differences due to the crystal packing form when complemented to the hexamer form (Fig. S6C). Moreover, apart from the observance of the distinctly different unit cell origin in the asymmetric unit, the differences in biologically relevant structures have also been supported by cross correlation analysis by GNM (Fig. S6A,B).

**Figure 2.**
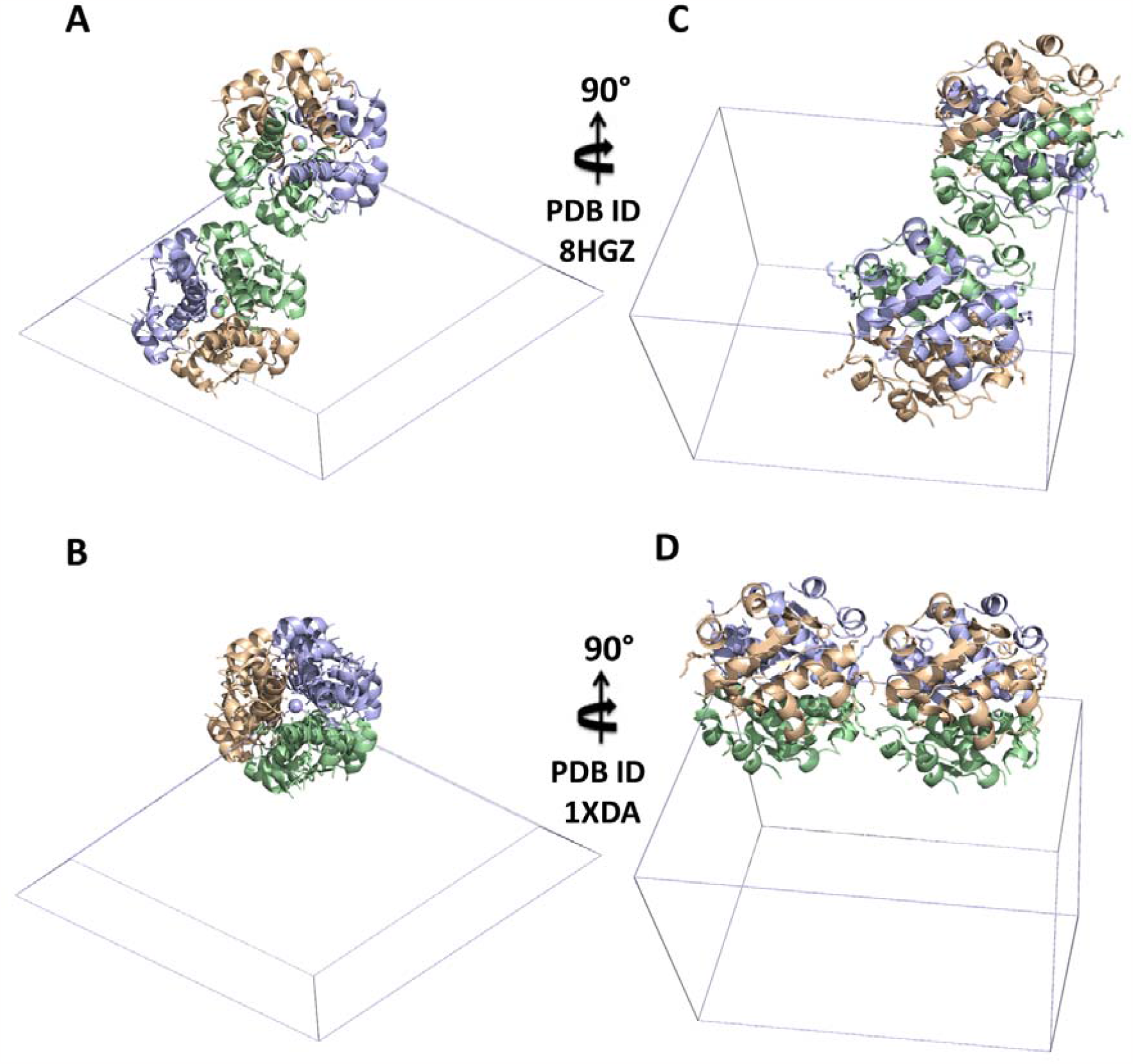
Comparison of the differences in unit cell origin of 8HGZ cryogenic structure and 1XDA cryogenic structure. (A) 8HGZ structure is presented in unit cell mode, also rotated 90 degrees (C). (B) 1XDA reference model is presented in unit cell mode, rotated 90 degrees (D) as well.

### Comparative GNM Analysis of Detemir in Two Different Crystal Forms

The GNM analysis is a minimalist coarse-grained method to reveal a protein’s dynamics [18]. Understanding the equilibrium fluctuations of the proteins is experimentally enabled by measuring the Debye-Waller/B-factor of each atom with X-ray crystallography [18]. Although the interpretation of crystallographic B-factors is complicated since they have represented the total of multiple sources of disorder, various analytical perspectives for decomposing molecular disorder into a hierarchical cluster of contributions can also provide an intuitive principle for quantitative structural-dynamics analysis [19]. Likewise, expected residue fluctuations can also be obtained by GNM in favorable agreement with experimentally measured fluctuations [18]. Thus, the relation between expected residue fluctuations obtained from GNM and thermal fluctuations/B-factors can be inspected in detail [18]. The fluctuation profile and the orientational cross correlations between pairs of nodes can also be obtained in agreement with experimental or adjustable parameters [18].

To observe if there is any difference that originated from unit cell differentiation in asymmetric unit cell, comparing their dynamics thoroughly, dimer of dimer form of the 1XDA and 8HGZ structures have been analyzed by GNM (Figure 3; Figure 4). Orientational cross-correlations between residue motions have been inspected and flexibility of the residues has been analyzed to describe the mean squared fluctuations. To investigate the communication in the residue networks, a cross-correlation map has been performed. In the cross-correlation heat-maps, blue-colored regions represent the uncorrelated motions, whereas red-colored regions represent the collective residue motions. Accordingly, AB-CD, and EF-GH have been observed as dimers with highly correlated motions in 1XDA structure (Figure 3A). In stark contrast, dimeric positive correlation in the 8HGZ detemir structure cannot be observed thoroughly due to extroverted monomers of the 1st dimer packing in the asymmetric unit cell (Figure 4A). Accordingly, the 1st dimer of the 8HGZ structure (AB-CD) has a weak correlation within each other (Figure 4A, bottom; Figure 4B, top), while the 2nd dimer of the 8HGZ structure (EF-GH) shows higher correlated motions compared to the first dimer (Figure S7A). In the 8HGZ crystal structure, residues between 1-40, 60-62, and 82-90 had at most25% correlation (Figure 4A bottom; Figure 4B). In addition, interchain residues 60-64 had at most 25% correlation with residues between 100-150, and 150-180. However, the interchain residue cross-correlation map of the 1XDA structure indicated that residue 120 had ∼25% correlation with residues 54-58 and residues 68-72, while residue 70 had ∼25% correlation with residues 104-108 and residues 118-122 (Figure 3A bottom; Figure 3B).

**Figure 3.**
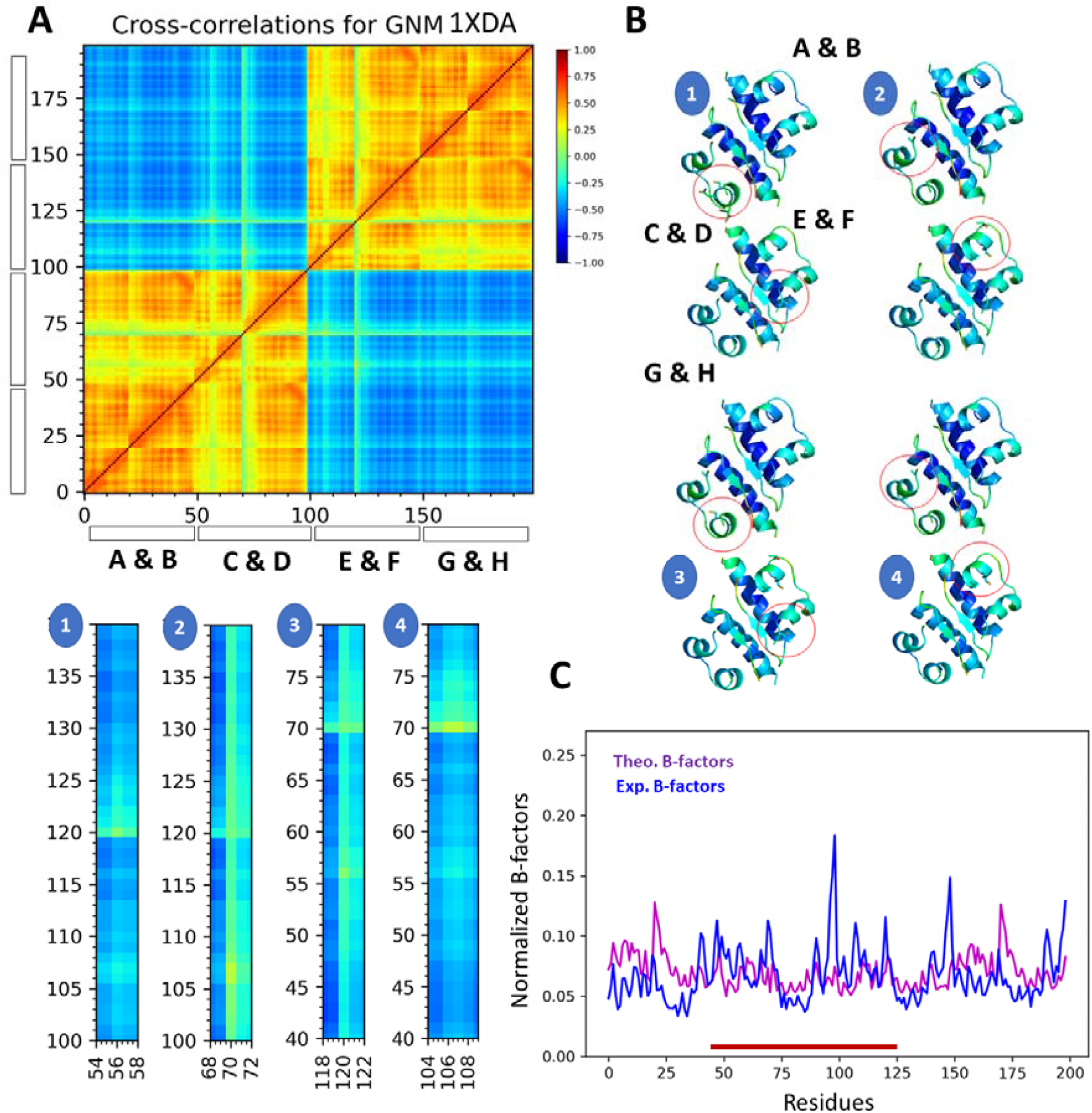
Gaussian Network Model (GNM) analysis results for 1XDA structure. (A) Crosscorrelation heat-map from overall GNM modes for 1XDA insulin structure and ∼25% correlated critical residues (54-58, 68-72, 104-108, 118-122, and 56, 70, 105-107,120) The residue pairs that move in the same direction is colored in red regions (Cijorient > 0); the residue pairs moving in opposite directions are colored in blue regions (Cijorient < 0); uncorrelated pairs are colored in green (Cijorient = 0.0; color-code bar on the right scale). (B) Cartoon representation of 1XDA structure colored in Bfactor. Critical residues corresponding to the heat map are numbered and emphasized in red circles. (C) Normalized B-factors of 1XDA are colored in blue, and theoretical B-factors calculated by GNM are colored in magenta. Critical residues corresponding to the heat map are emphasized in the red line.

**Figure 4.**
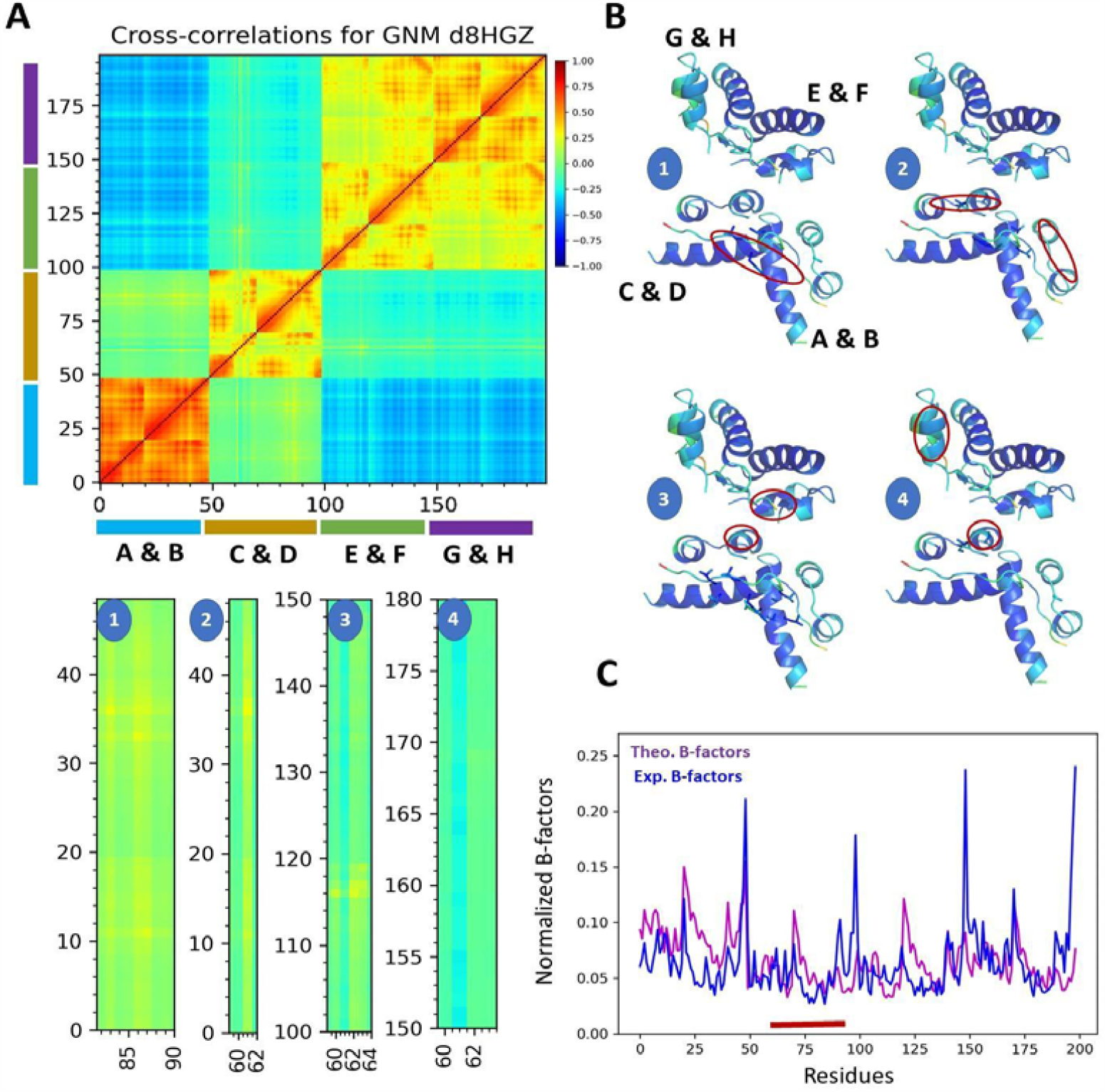
Gaussian Network Model (GNM) analysis results for 8HGZ structure. (A) Crosscorrelation heat-map from overall GNM modes for our structure and ∼25% correlated critical residues in AB-CD dimer. The residue pairs that move in the same direction are colored in red regions (Cijorient > 0); the residue pairs moving in opposite directions are colored in blue regions (Cijorient < 0); uncorrelated pairs are colored in green (Cijorient = 0.0; color-code bar on the right scale). (B) Cartoon representation of our structure colored in B-factor. Critical residues corresponding to the heat map are numbered and emphasized in red circles. (C) Normalized B-factors of the structure are colored in blue, and theoretical B-factors calculated by GNM are colored in magenta. Critical residues corresponding to the heat map are emphasized in red lines.

On the other hand, the first monomer (AB) displays at least 75% intramolecular correlation, followed by the last monomer (GH), which indicates higher (at least 50%) intramolecular correlation compared to monomer CD, (∼50%) and monomer EF (50%) (Figure 4A). A distinct correlation has been observed at the selected the same cut-off distance of 8Å; in the GNM of 1XDA structure, overall correlation with B-factors was 0.650 (Figure 3C), and 0.501 in the GNM of 1XDA structure (Figure 4C). Accordingly, contrary to the theoretical B-factor calculation, the experimental B-factor calculation is characterized by the highest fluctuation around 50th, 100th, 150th, and 200th residues in 8HGZ crystal structure (Figure 4C).

However, the B-factor analysis of the 1XDA structure around the 100th residue demonstrates higher fluctuation than other regions (Figure 3C). Remarkably, critical residues among the 175-198 and 165-170 are highly correlated with the 130-148 region in dimer EF-GH(Figure S7A). Especially, His 131 and Leu 132 in EF monomer is highly correlated with His 176 and Leu 177 (Figure S7C), supplemented by normalized b-factor fluctuation. Accordingly, when comparing the intra- and intermonomer residue motions of the 8HGZ structure, AB monomer displays at least 75% intramolecular correlation, the highest correlated chain among the others (Figure S8A, B). Interestingly, the C-terminal residues between 49-50 in chain B are highly correlated with the critical residues 73, 76-77, 81-82, 85, and 92-98 in chain D (Figure S8C, D). Similarly, the C-terminal residues 99 and 100 in chain D are highly correlated with around critical residues 123, 126-127, 130-131, 134, and 145-148 in chain F (Figure S9A, B). Likewise, the C-terminal residues 149 and 150 in chain F are highly correlated with the critical residues 173, 176-177, 180-181, 184-185, 187-188, and 195-198 in chain H (Figure S9C, D), meaning the intramonomer’ correlation pattern is quite similar in each chain (Figure S8; Figure S9). Collectively, comparative cross-correlation analysis for intra- and intercorrelation motion in AB, CD, EF, and GH monomers of 1XDA and 8HGZ displays a similar correlated pattern; however, intra- and intermolecular regions in 1XDA structure are highly-entirely correlated with each other compared to the corresponding regions in 8HGZ structure (Figure S10).

### Radiation damage comparison of the 8HGZ structure and 1XDA structure

Radiation damage of two structures was calculated using the RABDAM program [20]. B Damage values were performed using the total atomic isotropic B-factor values of determined atoms. The results are presented in kernel density plots in Figure S11. The highest B Damage value of 2.05 was observed on the Asn 18 C atom (541) in chain C of the 1XDA structure, while in the 8HGZ structure, the highest B Damage value (3.11) was observed on the Glu 154 O atom (1312) (Figure S11)

## Discussion

Insulin detemir is the first protein structure determined with an engineered fatty acid linkage/conjugation [11]. Even though lipid modification of the proteins occurs routinely in nature, engineering strategies assist in understanding novel possibilities in oligomeric interactions, providing an enabling biotechnological tool for practical applications in drug therapy [11]. While numerous insulin structures are studied in dimer form, the presence of the fatty acid chain in the insulin paves the way to inspect the insulin in two-dimer form [7,9,10]. This provides the understanding of the dynamic profile of the molecule from an oligomeric perspective. Thus, the assembly of subunits interacting through the covalent and noncovalent bonds defines their structure-function correlations [21]. Ideally, phenolic ligands and zinc atoms retain the hexameric form of insulin [22]. The rationale for the long-acting form of the insulin detemir is characterized by keeping the molecule in the dimer form of hexamers compared to other protracted insulins [22]. Thus, after injection of the detemir subcutaneously, phenolic ligands and fatty acid groups attached to insulin rapidly diffuse into the subcutaneous area, establishing the hexamer:dihexamer equilibrium [22]. The accumulated forms slowly dissociate into dimers and monomers, after which the myristic acid groups on insulin permit self-assembly, prolonging its action [11, 22] (Figure S2).

The oligomeric state of insulin detemir may further contribute to the understanding of monomeric-dimeric correlated motion by demonstrating comparable properties to 1XDA structure. In this work, we examined the interaction and correlation of each monomer and dimer state of the oligomeric insulin detemir whose crystal structure determined in a novel crystal packing form in asymmetric unit cell (Figure 1; Figure S3; Figure S4; Table 1). Notice that the presented structure here has a distinct unit cell origin compared to the 1XDA structure (Figure 2; Figure S5). In our structure, two dimers of the 1XDA indicate that each dimer has an introverted state (Figure 3B), while the 8HGZ have an extroverted dimer form in the unit cell (Figure 4B). Namely, the “dimer of dimer” form of 8HGZ displays minor differences due to the crystal packing form when complemented to its dihexamer form (Figure S6).

Accordingly, in GNM analysis, the second dimer of the presented structure has a higher correlation with each other compared to its first dimer (Figure S7; Figure 4). Moreover, the C-terminal residues, Pro and Lys, in chains B, D, F, and H displayed correlated motions with the critical regions in chains D, F, and H, respectively (Figure 5; Figure S8; Figure S9).

**Figure 5.**
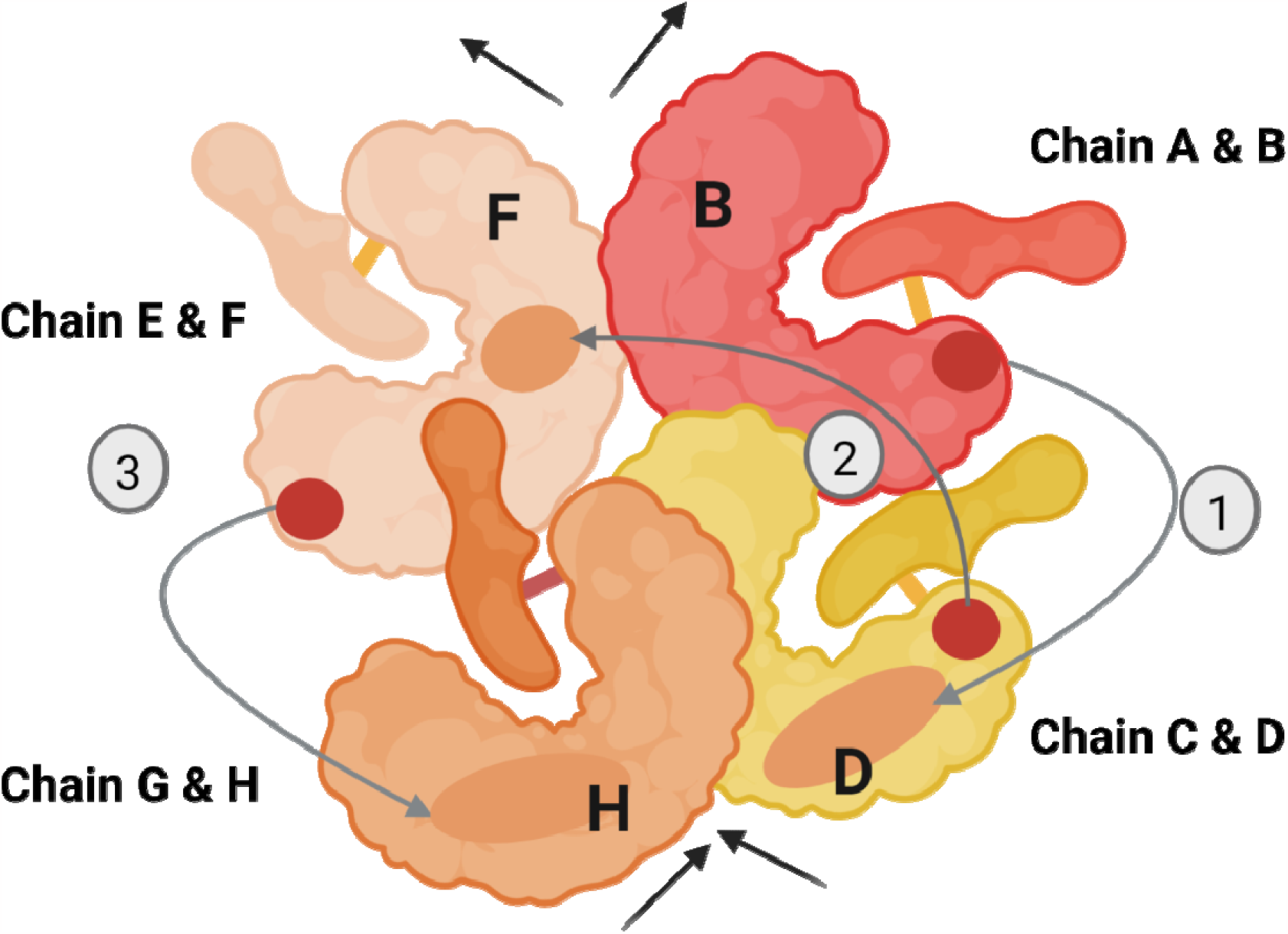
Pictorial representation of monomeric-dimeric correlation of the 8HGZ cryogenic structure. According to cross-correlation with GNM analysis of 8HGZ structure, the C-terminal Pro and Lys residues of chain B are highly correlated with chain D (1); the C-terminal Pro and Lys residues of chain D are highly correlated with chain F (2), and the C-terminal Pro and Lys residues of chain F are correlated with chain H (3). 8HGZ structure composed of two dimers or four monomers; compared to 1XDA structure, AB-CD (1st dimer) are extroverted formswhereas EF-GH (2nd dimer) are slightly introverted formsthan 1st dimer. Based on the intrachain’ correlation, AB monomer has at least a 75% correlation with each other (colored in red); CD monomer and EF monomer correlate in intrachain up to 50% (colored in yellow, and wheat); GH monomer has at least 50% correlation with each other (colored in orange).

Interestingly, the first monomer of insulin shows at least 75% intramonomer correlation with almost all residues, followed by the last monomer of the insulin, which shows higher intramolecular correlation compared to the other chains (Figure 4A). At neutral pH, detemir exists as a dimer of dimer form in the asymmetric unit [11]. The best estimated dissociation rate of an insulin dimer into two monomers in water is 0.4 μs-1 [23]. Our best crystallization condition of the insulin detemir has occurred at pH 9. Considering that the dissociation of insulin occurs more effective at higher pH values (pH ≥ 8) and reaches the maximum by pH 10 [24], the minor distinct conformational arrangement of the presented structure’s critical residues compared to 1XDA structure might be related to the slow dissociation of the structure during crystallization. In a study conducted to observe the conformational changes of insulin crystals in the pH range of 7-11, observed that the conformational changes were closely related to the pH value, which is compatible with our minor conformational changes in symmetry mates’ analysis obtained from 1XDA vs 8HGZ dihexamer form. [25]. Correlated motion of each subunit represented in Figure 5 may confirm the causal interrelationship of the oligomeric insulin’ dissociation (Figure S8; Figure S9). Additionally, comparative cross-correlation of 1XDA and 8HGZ structures has demonstrated that the 1XDA monomer-dimer correlation is entirely +75-100% rate (Figure S10). However, the corresponding regions in 8HGZ have indicated a weak correlation between monomers and dimers, suggesting a loosely coupled arrangement following the slow dissociation of the crystal structure occurred due to elevated pH value. Moreover, observing increased fluctuations in relevant Pro and Lys residues in normalized B-factors (Figure 4C) might confirm the cooperative motion originated from the alternative conformation of the oligomeric insulin (Figure 5). Ideally, insulin lispro has been introduced as a rapid-acting insulin analog, reversing the proline at position B28 and lysine at position B29 [26]. This substitution causes a conformational shift in the C terminal of the chain, inhibiting the ability of the insulin monomers to form dimers sterically [26]. Therefore, the dissociation constant of the dimer is reduced by 300-fold compared to regular insulin [26]. Previous studies suggest that proline has an inhibitory effect on the protein fibrillation process, promoting the retention of protein conformation [27,28]. Considering the cross-correlation of C-terminal residues with neighboring monomers (Figure S8; Figure S9), it could be argued that the detemir insulin demonstrates oligomeric cooperativity via the Pro and Lys residues (Pro 49, Lys 50; Pro 99, Lys 100; Pro 149, Lys 150). On the other hand, C-terminal lysine provides a platform for covalent binding of the structure with the fatty acid, allowing for the long-acting profile of the insulin [11]. Together with the GNM results, these could mean that the dissociation of the oligomeric insulin begins with the Brownian motions and fluctuations of the myristic acid.

To determine and validate the accuracy of the monomeric-dimeric correlated motion of the protracted insulins, coarse-grained normal mode analysis offers a robust dynamic profile for future studies. Furthermore, when exposed to global radiation damage, diffraction patterns can be disrupted, causing an increase in both unit-cell volume and mosaicity. This increase in volume can result in non-isomorphism, which makes determining the structure more challenging. Therefore, considering the radiation damage of the 8HGZ structure compared to the 1XDA structure (Figure S11), the next step is to understand the details of this dynamic profile by a time-resolved ambient temperature radiation-damage free Serial Femtosecond X-ray (SFX) crystallographic studies supplemented with diffuse X-ray scattering.

## Materials & Method

### Sample preparation and crystallization

The detemir sold as brand name Levemir® was crystallized by employing sittingdrop microbatch vapor diffusion screening technique under oil using 72-well Terasaki crystallization plates as explained in Ertem et al. 2022 [29]. Briefly, the detemir solution was mixed with (1:1 ratio, v/v) ∼3500 commercially available sparse matrix crystallization screening conditions at ambient temperature for crystallization. Each well containing 0.83 μl protein and crystallization condition was sealed with 16.6 μL of paraffin oil (Cat#ZS.100510.5000, ZAG Kimya, Türkiye) and stored at room temperature until crystal harvesting. Terasaki plates were checked under a compound light microscope for crystal formation and growth. Large crystals were obtained within 48 hours. The best crystals were grown in a buffer containing 0.2 M sodium acetate trihydrate, 0.1 M TRIS hydrochloride pH 9.0.

### Crystal harvesting

The crystals were harvested from the Terasaki crystallization plates using MiTeGen and micro loop sample pins mounted to a magnetic wand under the compound light. microscope. The harvested crystals were immediately flash-frozen by plunging in the liquid nitrogen and placed in a previously cryo-cooled sample puck (Cat#M-CP-111-021, MiTeGen, USA). The filled puck within the liquid nitrogen was transferred to the dewar of the Turkish DeLight for diffraction data collection [17].

### Data collection and processing

Diffraction data was collected by using Rigaku’s XtaLAB Synergy Flow XRD equipped with CrysAlisPro software (Agilent, 2014) as described in Atalay et al. 2022 [17]. Before the data collection, the system was cooled to 100 K by adjusting Oxford Cryosystems’s Cryostream 800 Plus. After that, the dewar was filled with liquid nitrogen and a loaded sample puck was placed into the cryo sample storage dewar installed at the Turkish DeLight. The robotic auto sample changer (UR3) mounted the MiTeGen sample pin inside the sample storage dewar to the Intelligent Goniometer Head (IGH) by the 6-axis robotic arm. The crystal was centered using the automatic centering feature of the CrysAlisPro software to the X-ray interaction position. For the data collection, the PhotonJet-R X-ray generator with Cu X-ray source was operated at full-power of 40 kV and 30 mA, and the beam intensity was set to 10% by using piezo slit system to minimize the cross-fire and overlap of the Bragg’s reflections. The diffraction data, which contains 1800 frames in total, is collected to 1.7 Å resolution at a 45 mm detector distance, 0.2-degree scan width, and 1-second exposure time. The data was processed by the automatic data reduction feature of CrysAlisPro. When data reduction is completed, the output files, which contain both unmerged and unscaled files (*.rrpprof format) converted to the structure factor file (*.mtz format) through the *.hkl file. This step is performed by integrating CCP4 crystallography suite [30] into CrysAlisPro software.

### Structure determination and refinement

The high-resolution crystal structure of detemir is determined to 1.7 Å resolution at cryogenic temperature in space group R3:H. Molecular replacement is performed by using the automated molecular replacement program PHASER [31] implemented in PHENIX software package [32] by using the previous detemir X-ray crystal structure at cryogenic temperature as a search model [11] (PDB ID: 1XDA). Coordinates of the 1XDA were used for the initial rigid-body refinement with the PHENIX software package. After simulated-annealing refinement, individual coordinates and Translation/Libration/Screw (TLS) parameters were refined. The potential positions of altered side chains and water molecules were checked in COOT [33], and positions with strong difference densities were retained. Missing water molecules were manually added, or water molecules located outside of significant electron density were removed. The Ramachandran statistics for detemir structure (most favored / additionally allowed/disallowed) are 100% / 0.0% / 0.0%, respectively. The structure refinement statistics are summarized in Table 1. All X-ray crystal structure figures were generated with PyMOL [34].

### GNM analysis

The 1XDA and 8HGZ structures were analyzed by Normal Mode Analysis using ProDy [18]. Orientational cross-correlation map was defined with all Cα atoms of the proteins (residues 0-200). Myristic acids, phenols, zinc, and chloride atoms were excluded from the analysis. The cut-off distance of 8 Å was preferred in both structures to assume pairwise interactions, and the default spring constant was defined as 1.0 for both models. 1883 and 1673 atoms were parsed, and 198 non-zero modes were calculated with GNM for 8HGZ and 1XDA structures, respectively. The experimental B-factors were compared with the theoretical fluctuations calculated as well as orientational cross-correlations between residue fluctuations were calculated over all GNM modes. The differences in cross-correlations between 8HGZ and 1XDA structures were calculated at determined sections. Accordingly, intra-chain cross-correlations of each monomer in 8HGZ were compared to the intra-chain cross-correlations of each chain in the 1XDA reference model; and the interchain cross-correlations between all chains were calculated similarly, presenting as heat-maps.

### Normalization of B-factor

B-factor normalization involves transforming experimental B-factors to define their distribution in terms of expected value and variance. This allows for comparing B-factors across different datasets while eliminating prominent influences that may skew the data. To carry out this normalization, the pdb library transfers the normalized data into a temporary record set containing B’-factors per residue. The first algorithm for B-factor normalization was proposed by Karplus and Schulz, which related the experimental B-factor of a given residue to the arithmetic mean of all B-factors in a structure [35]

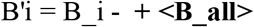

where B’i is the normalized B-factor of residue i, B_i is the experimental B-factor of residue i, is the average B-factor for all residues in the structure, and <B_all> is the average B-factor for all atoms in the structure.

## Conclusions

Here we present the detemir insulin structure at 1.7Å resolution, displaying distinct unit cell origin in the asymmetric unit. Considering comparative structural and computational analysis with 8HGZ and 1XDA structures, this study suggests that elevated pH can induce an intermediate state in the dissociation pathway of a protracted insulin analog. The distinct correlations especially induced at Lys and Pro residues in the C-terminus of the B, D, F, and H chains, can offer a clue about the insulin detemir’s hexamer:dihexamer equilibrium. Coarse-grained normal-mode analysis to determine the accuracy of monomeric-dimeric correlated motion of long-acting insulins provides a robust dynamic profile for future studies.

## Acknowledgment/Disclaimers/Conflict of interest

Authors would like to dedicate this manuscript to the memory of Dr. Albert E. Dahlberg and Dr. Nizar Turker. The authors gratefully acknowledge the use of the services and facilities of the Koç University Isbank Infectious Disease Center (KUISCID). H.D. acknowledges support from NSF Science and Technology Center grant NSF-1231306 (Biology with X-ray Lasers, BioXFEL). A.K. acknowledges support from the Scientific and Technological Research Council of Türkiye (TÜBİTAK, 2218 - National Postdoctoral Research Fellowship Program under project number 118C476 and 122F301). This publication has been produced benefiting from the 2232 International Fellowship for Outstanding Researchers Program, the 1001 Scientific and Technological Research Projects Funding Program of the, 2244 Industry Academia Partnership Researcch Project Funding Program and 2236 CoCirculation2 Program of TÜBİTAK (Project Nos. 118C270, 120Z520, 119C132 and 121C063). However, the entire responsibility of the publication belongs to the authors of the publication. The financial support received from TÜBİTAK does not mean that the content of the publication is approved in a scientific sense by TÜBİTAK. Coordinates of the detemir structure have been deposited in the Protein Data Bank under accession code 8HGZ.

---

**Supplementary Figure 1.**
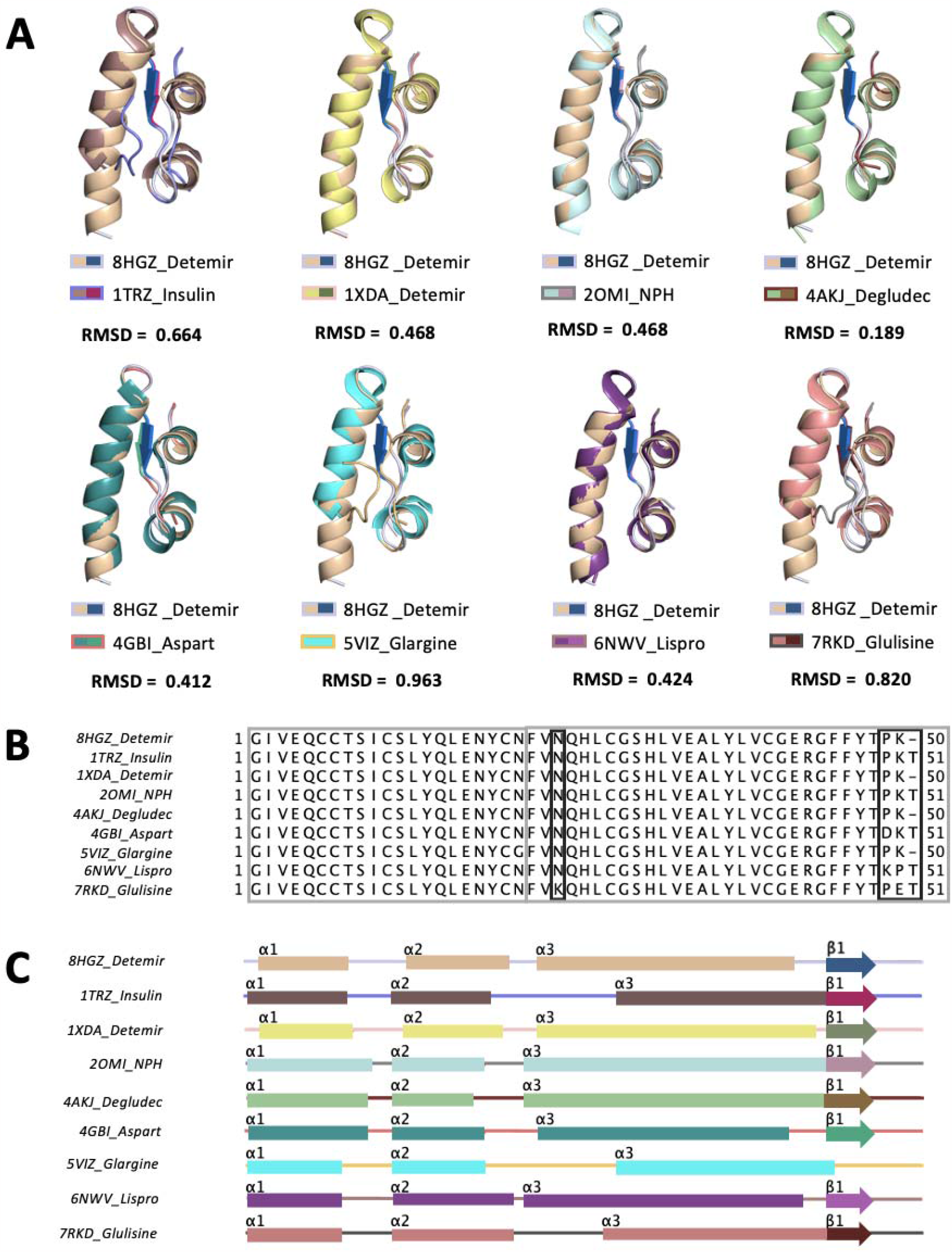
Alignment of detemir structure at cryogenic temperature with insulin analogs. (A) The monomer form of detemir (PDB ID: 8HGZ) is superposed with insulin analogs in *PyMOL*. Each structure is colored based on a secondary structure (helix, beta sheet and loop) which is detailly represented in Panel C. (B) Sequence alignment is performed using Jalview, and (C) secondary structures are indicated with different colors for each insulin homologs based on PDB IDs. (PDB ID: 1TRZ native human insulin; PDB ID: 1XDA insulin detemir; PDB ID: 2OMI NPH insulin analog; PDB ID: 4AKJ insulin degludec; PDB ID: 4GBI insulin aspart; PDB ID: 5VIZ insulin glargine; PDB ID: 6NWV; insulin lispro; PDB ID: 7RKD insulin glulisine.) Coordinates of the detemir structure have been deposited in the Protein Data Bank under accession code 8HGZ.

**Supplementary Figure 2:**
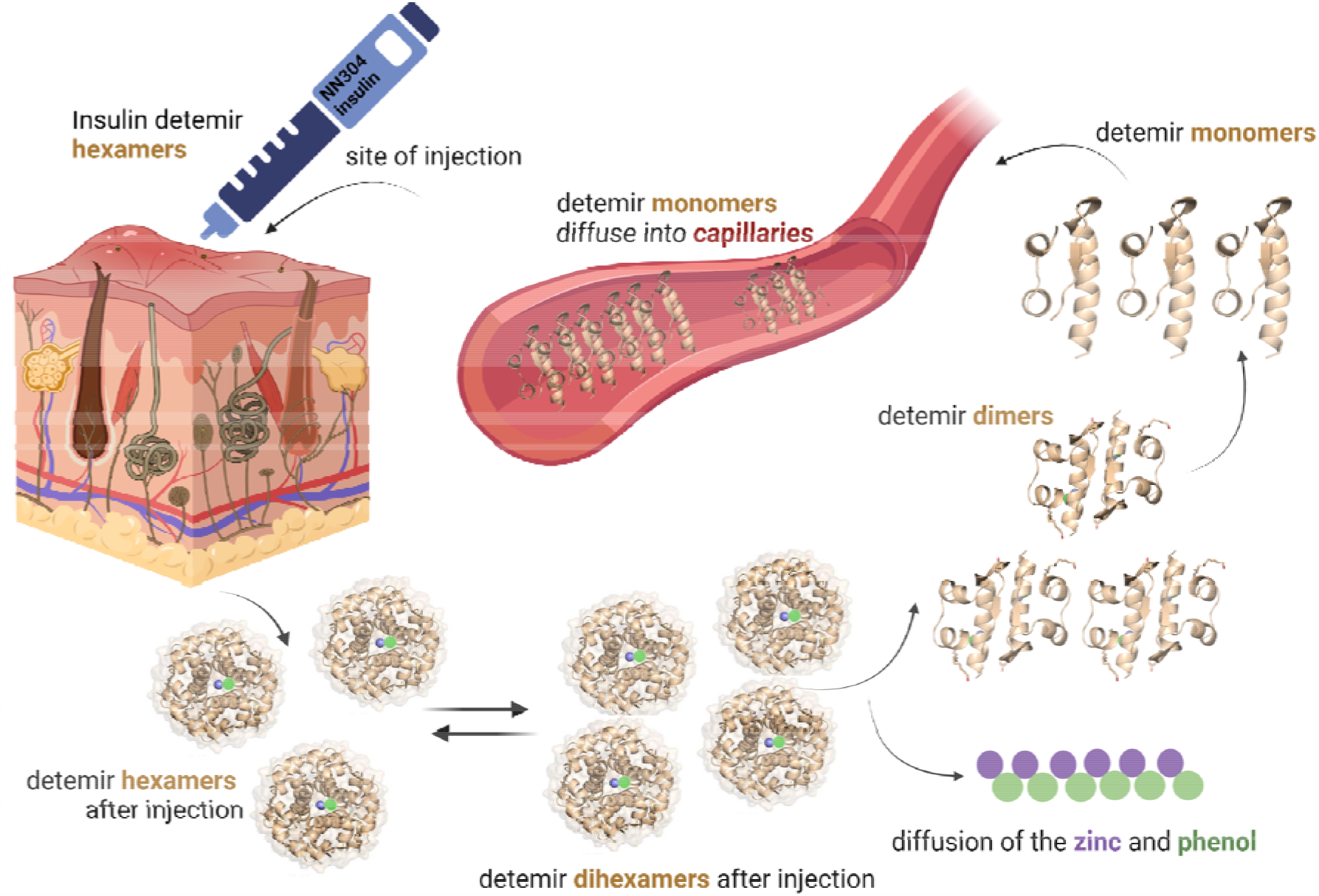
Summary of the action mechanism of the insulin detemir. Once insulin is injected subcutaneously hexamer or dihexamers diffuse and dissociate slowly in the blood. Insulin detemir dihexamers, hexamers, dimers, and monomers are colored in wheat, zinc and phenol are shown as circles and colored in purple and green, respectively.

**Supplementary Figure 3.**
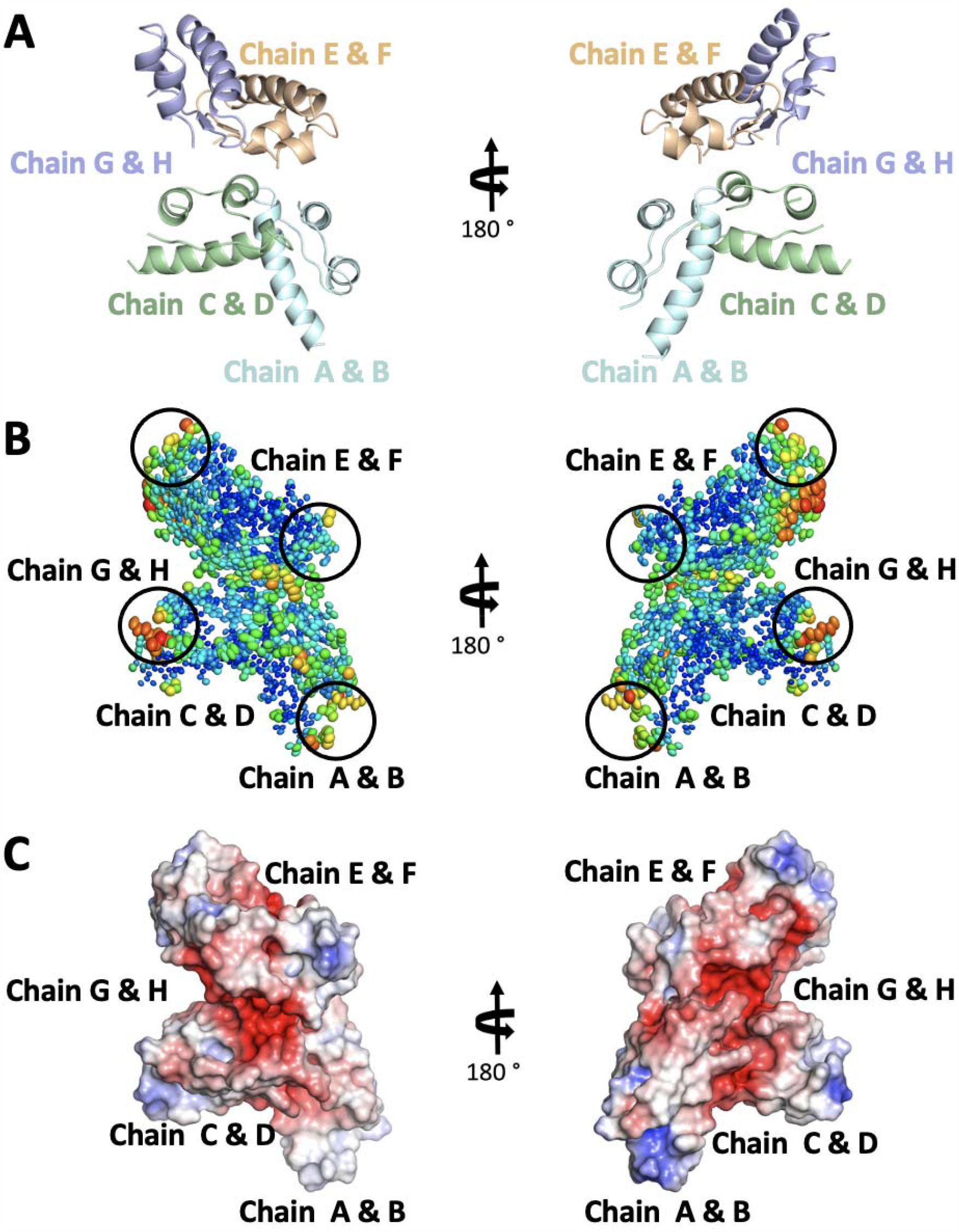
Ellipsoid and electrostatic representation of 8HGZ structure at cryogenic temperature. (A) Each monomer of detemir is indicated with a distinct color. (B) Detemir is shown with the ellipsoid presentation to highlight the stable/flexible regions shown in warmer colors. The binding site of myristic acid (MYR) is indicated with circles.. (C) Electrostatics is generated to show the surface charges on detemir by using the APBS electrostatic plug in of *PyMOL*.

**Supplementary Figure 4.**
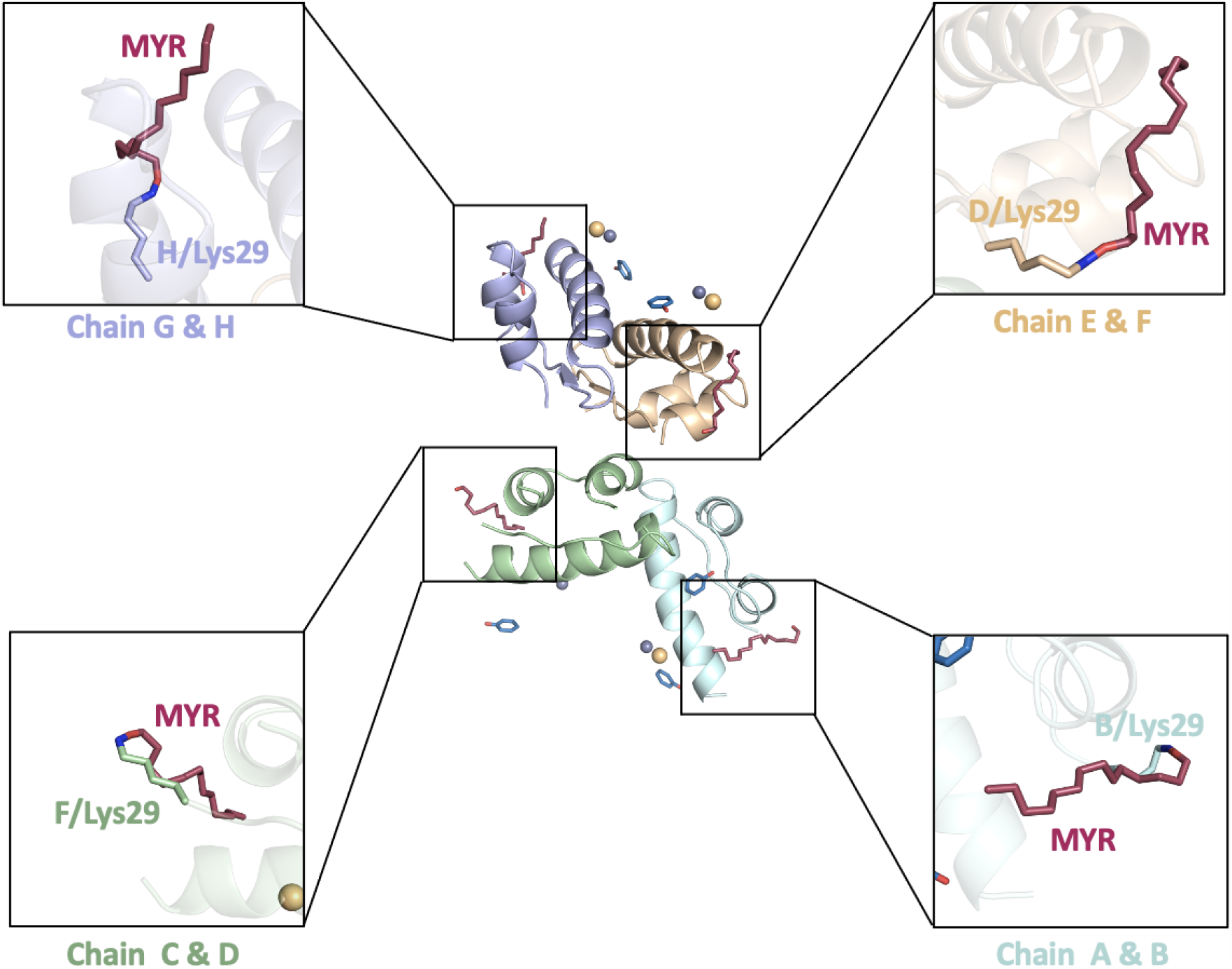
Covalent bond representation between myristic acid (MYR) and detemir. The binding of each MYR to detemir is shown with stick representation. The residues are highlighted based on atom representation to indicate the covalent bond with MYR.

**Supplementary Figure 5.**
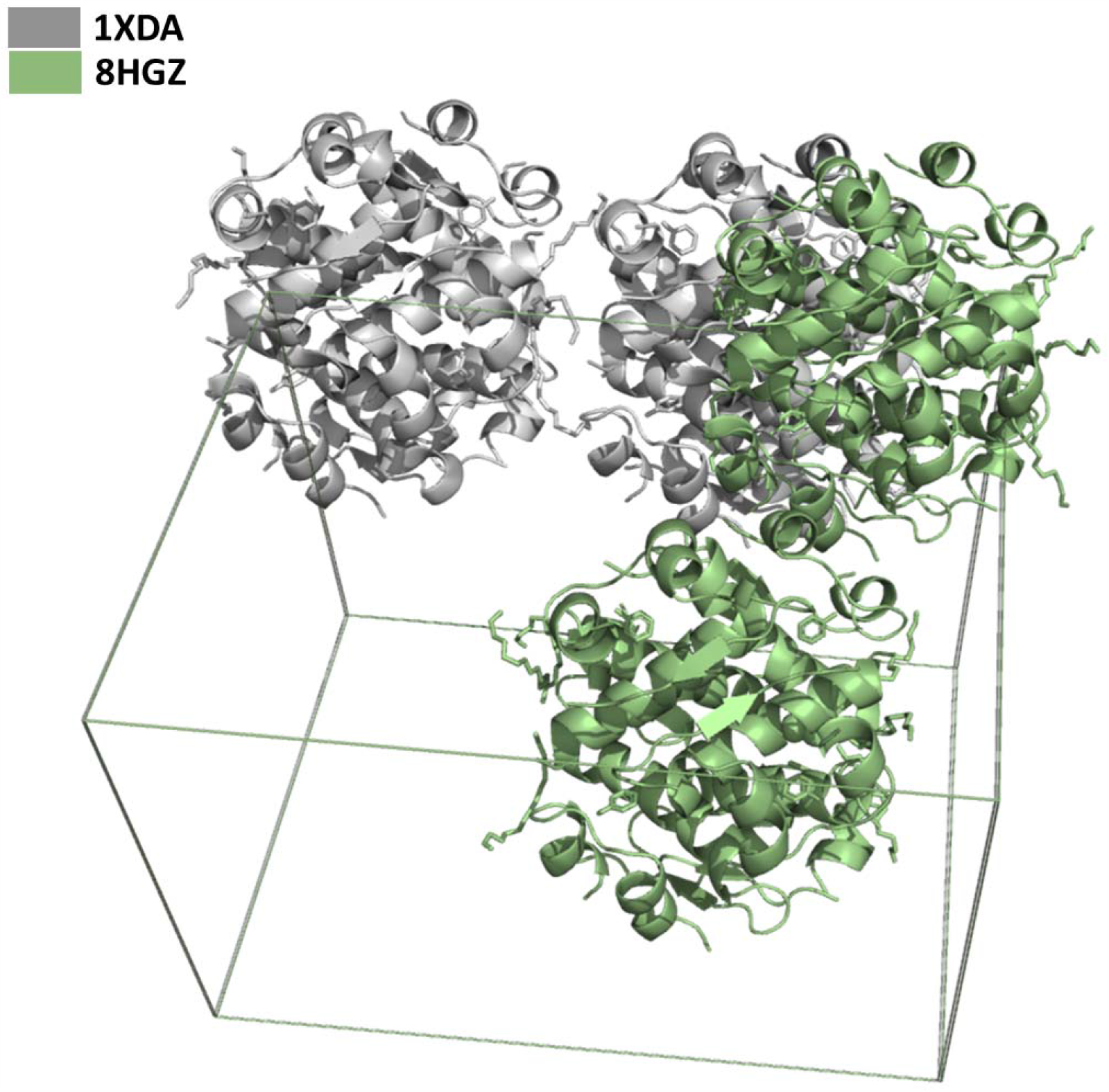
Superimposed two detemir structures in unit cell mode. 8HGZ structure is colored in green, while the 1XDA structure is colored in gray.

**Supplementary Figure 6.**
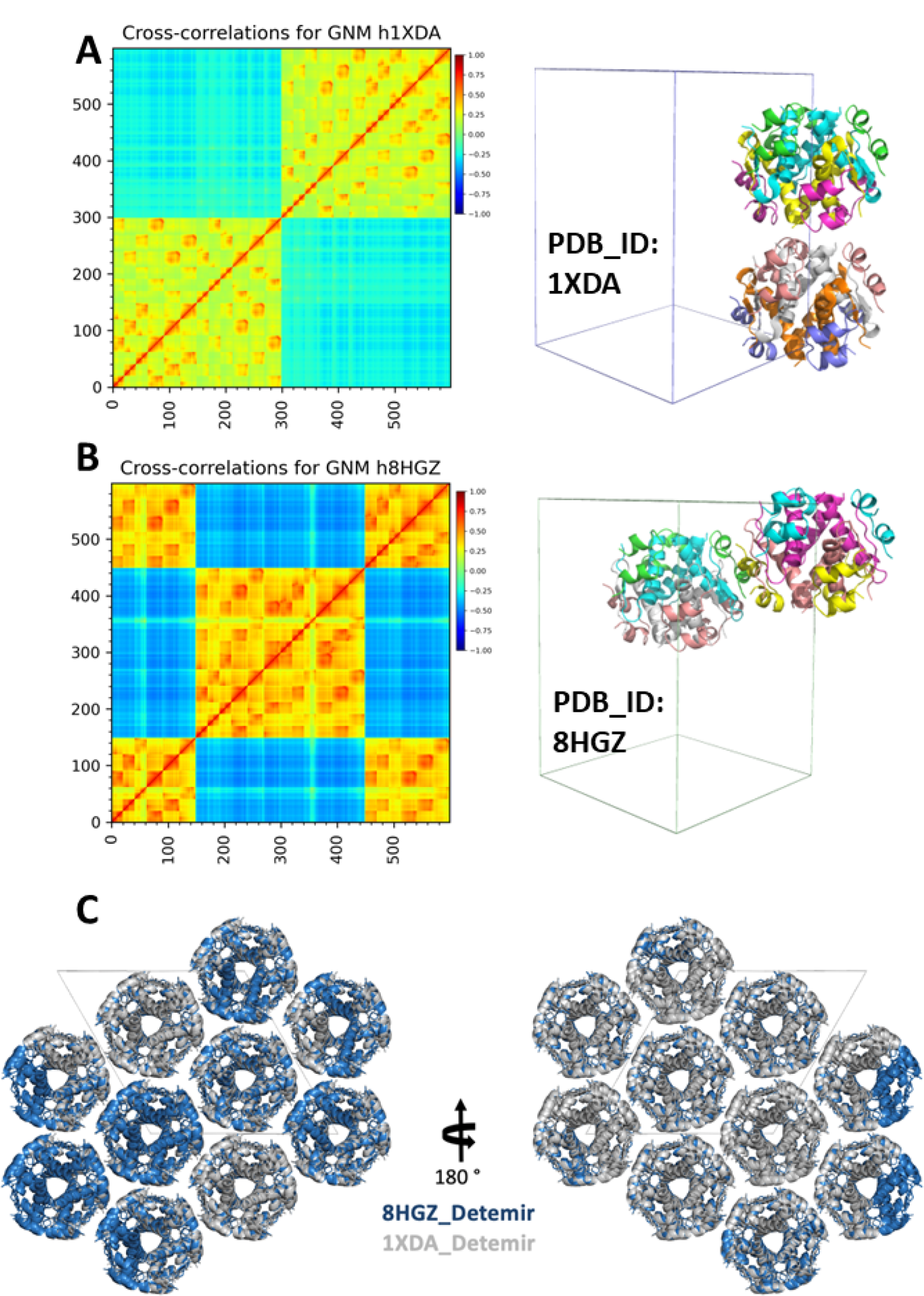
Representation the dihexamer conformation of the 8HGZ and 1XDA structures. (A) Cross-correlation analysis by GNM of the 1XDA’ dihexamer conformation and its cartoon representation in unit cell mode. (B) Cross-correlation analysis by GNM of the 8HGZ’ dihexamer conformation and its cartoon representation in unit cell mode. The cut-off distance of 8 Å was preferred in both structures to assume pairwise interactions. (C) Crystal packing representation of 1XDA (gray) and 8HGZ (skyblue) structures.

**Supplementary Figure 7.**
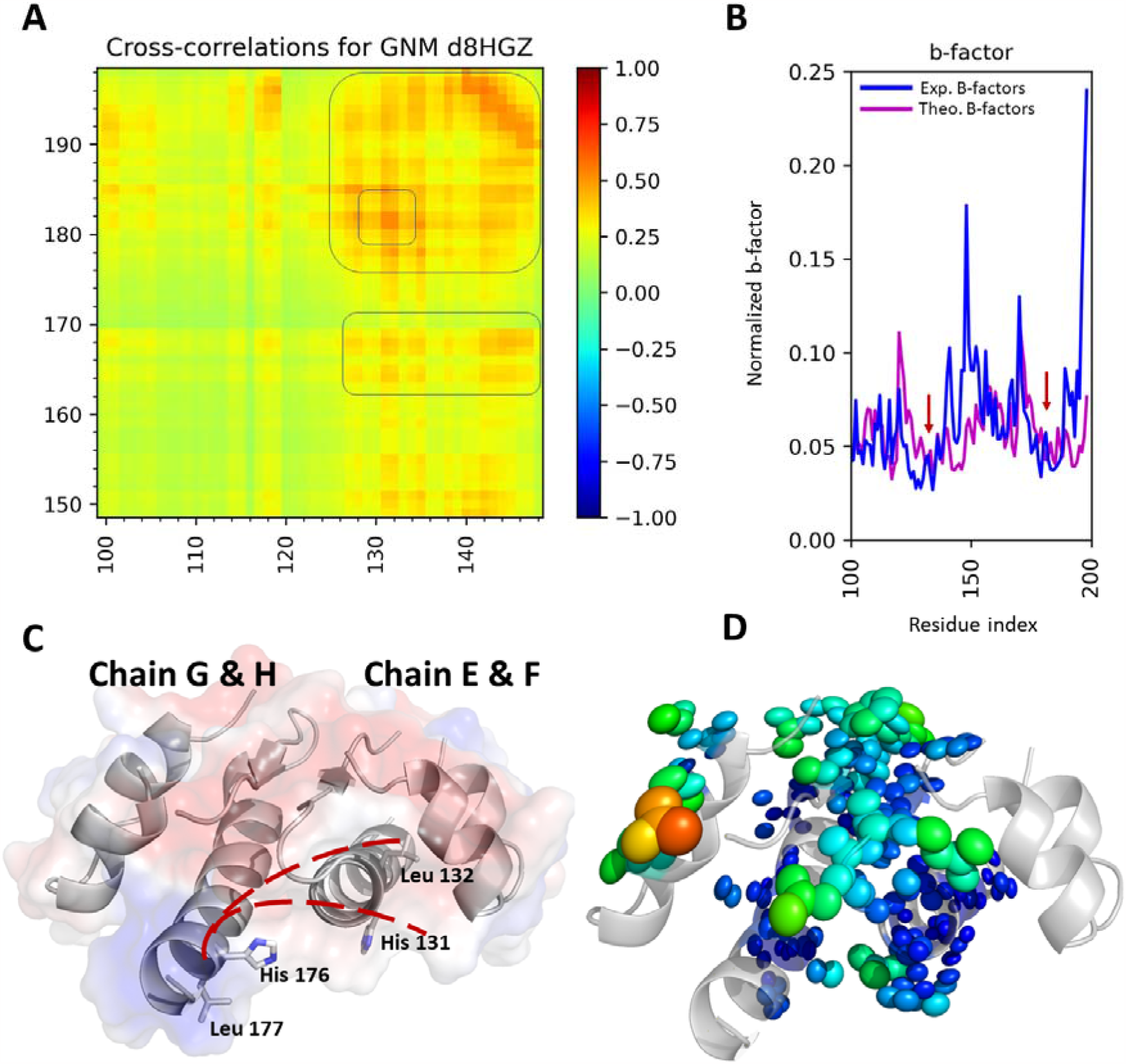
Gaussian Network Model (GNM) analysis results for the 2nd dimer of the 8HGZ structure. The 2nd dimer EF-GH) is more correlated than the first dimer (AB-CD; Fig.4A, bottom). The sliced model representation of the 2nd dimer and corresponding residues of the normalized b-factor plot has been calculated with GNM (A,B), supplemented with cartoon,electrostatic surface mode and ellipsoid representation to show the correlated motion of the dimer (C,D).

**Supplementary Figure 8.**
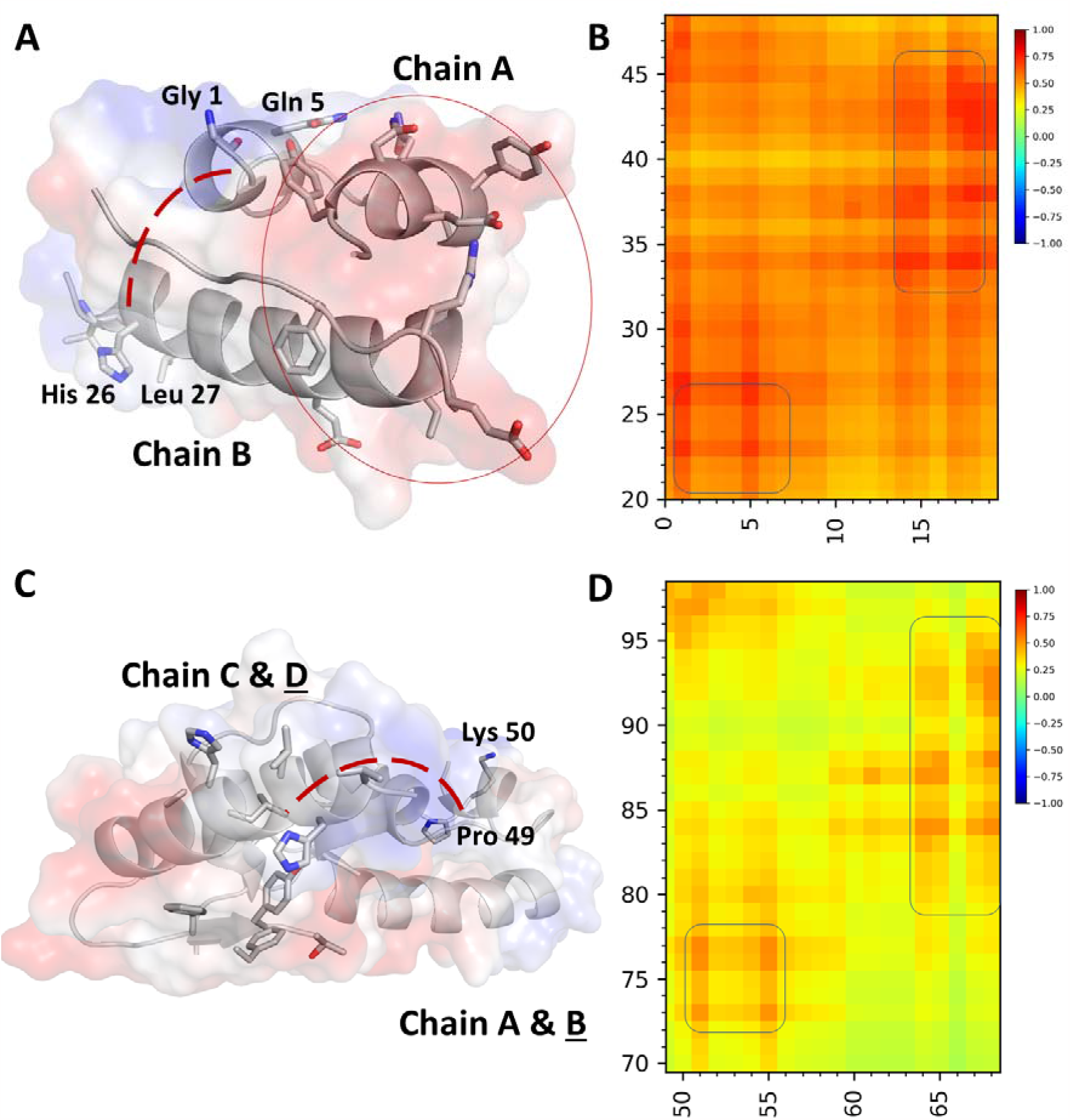
Gaussian Network Model (GNM) analysis results for intra- and intercorrelation motion in AB-CD dimer. The sliced model representation of chain A&B and chain C&D have been calculated with GNM (B, D), which is supplemented with cartoon and electrostatic surface mode to show the correlated motion of these chains (A, C). The C-terminal residues in chain B are highly correlated with chain D. Moving direction is emphasized by the red dashed line.

**Supplementary Figure 9.**
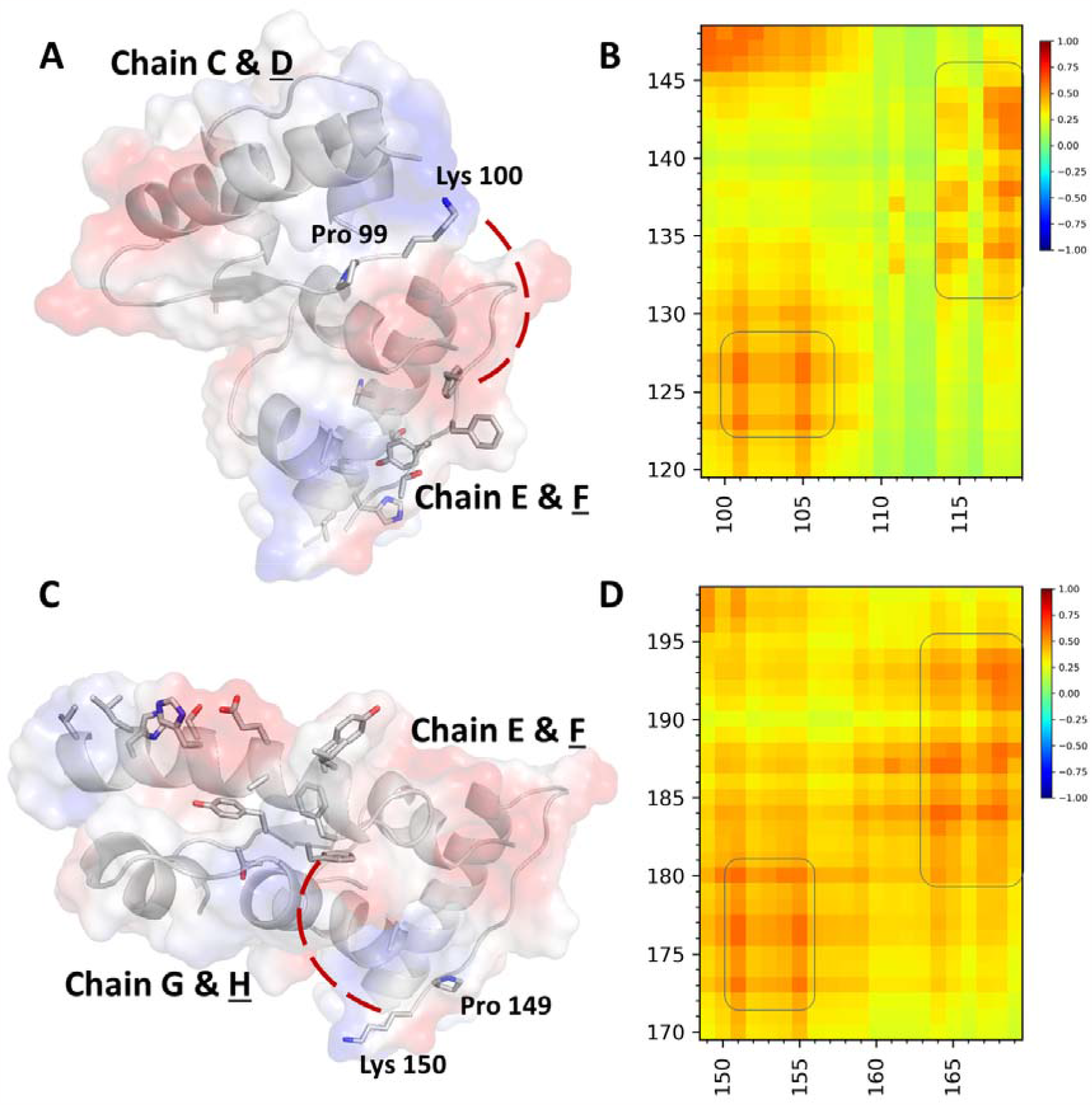
Gaussian Network Model (GNM) analysis results for intra- and intercorrelation motion in CD, EF, and GH monomers. The sliced model representation of EF and GH monomers have been calculated with GNM (B, D), which is supplemented with cartoon and electrostatic surface mode to show the correlated motion of these chains (A, C). The C-terminal residues in chain D are highly correlated with chain F; the last residues of chain F are highly correlated with chain H. Moving direction is emphasized by the red dashed line.

**Supplementary Figure 10.**
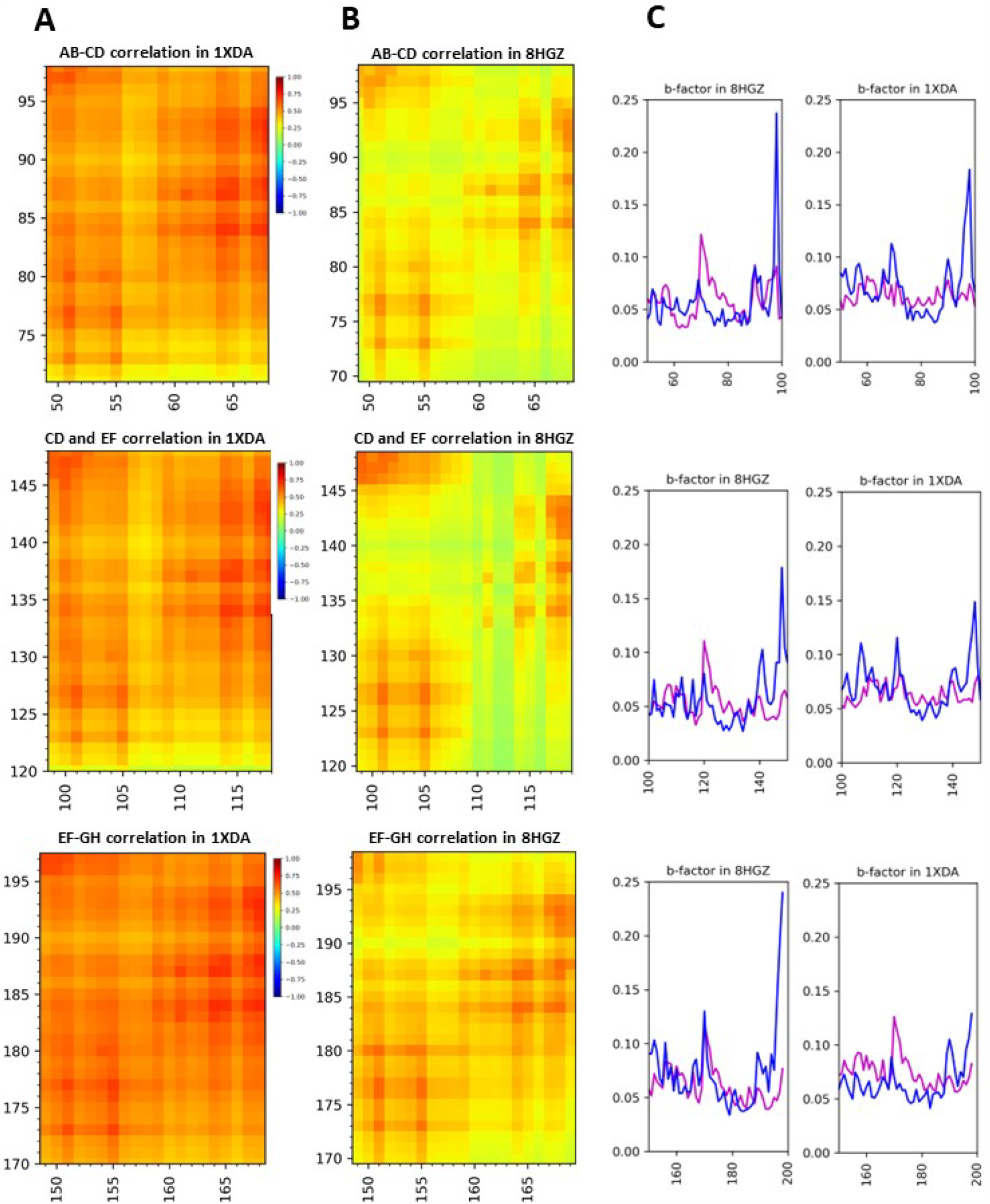
Comparative Gaussian Network Model (GNM) analysis results for intra- and intercorrelation motion in AB, CD, EF, and GH monomers of 1XDA and 8HGZ structures. (A) Intra- and intercorrelation motion of 1XDA’ monomers is highly correlated compared to (B) 8HGZ’ monomers. (C) Comparative normalized b-factor analysis of corresponding residue index in 1XDA and 8HGZ monomers. Magenta: Theorotical b-factor, Blue: Experimental b-factor.

**Supplementary Figure 11.**
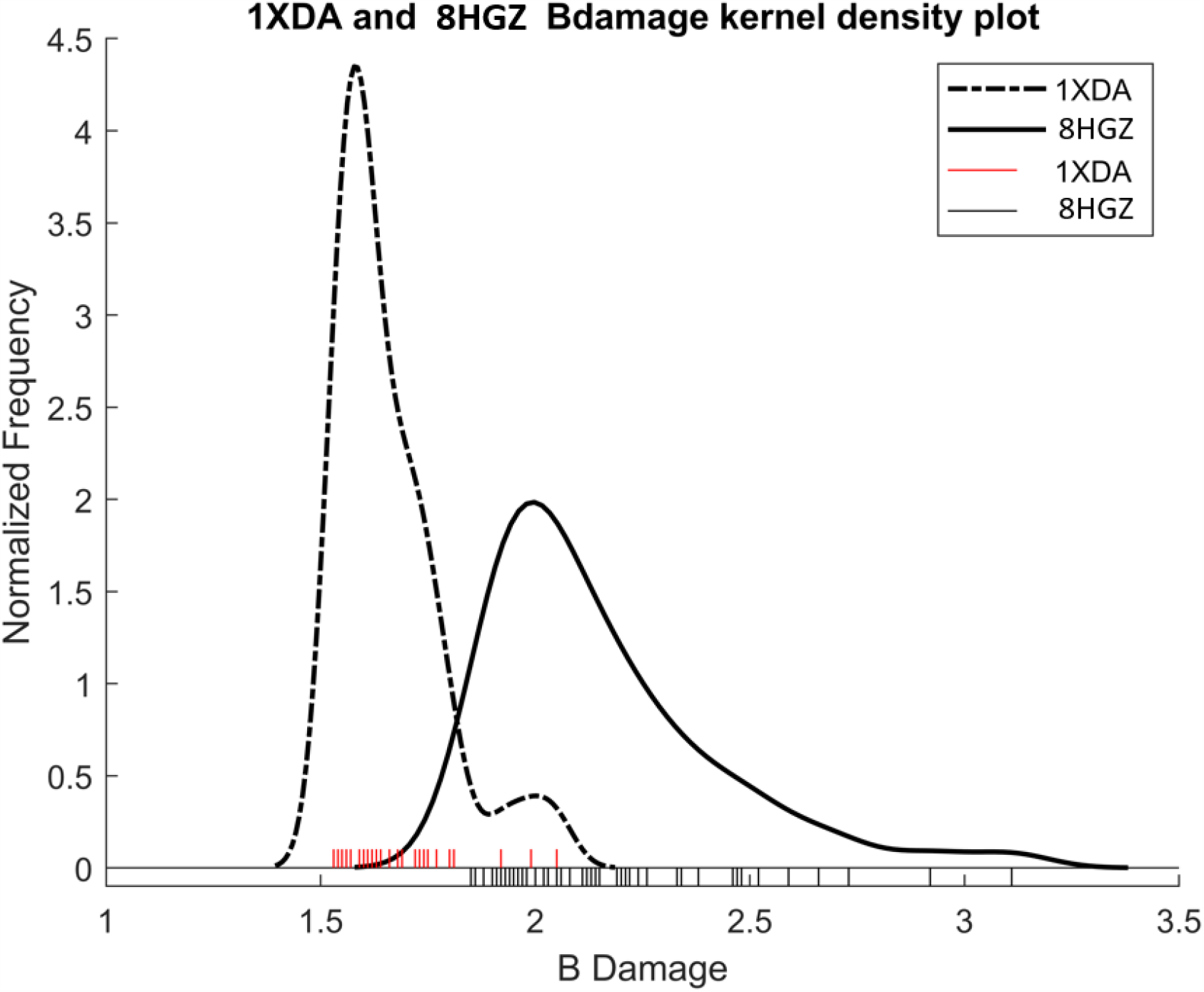
Radiation damage values were calculated using RABDAM software. Comparison of the BDamage distribution plot of the 8HGZ and 1XDA structures. The highest BDamage value of 3.11 was observed on the Glu 154 O atom (1312) of the 8HGZ structure, while in 1XDA structure, the highest BDamage value (2.05) was observed on the Asn 18 C atom (541) in chain C.

